# RNA editing is a molecular clock in unmodified human cells

**DOI:** 10.1101/2024.11.18.624170

**Authors:** Ali E. Ghareeb, James Bayne, Aaron Z. Wagen, Alya Masoud Abdelhafid, David Miller, Laura Cubitt, Laween Meran, Ciaran Hill, Adam P. Cribbs, Mina Ryten, Sonia Gandhi, Eamonn Gaffney, Mark Coles, George Young, Samuel G. Rodriques

**Author notes:** These authors contributed equally to this work (Ali E. Ghareeb, James Bayne, Aaron Z. Wagen).

## Abstract

Despite major advances in spatial RNA sequencing, the ability to extract temporal information in RNA sequencing experiments is still limited. Here, we describe Transcriptome Timestamping (T2), a system which harnesses naturally occurring A-to-I editing of RNA transcripts in unmodified human cells to infer transcriptional history. T2 provides age estimates for individual RNA transcripts, and serves as an endogenous molecular recorder, differentiating between complex transcriptional programs. We show that T2 can identify transient and transitional transcriptional programs in primary differentiating monocytes that are not apparent from gene expression analysis alone, including a regulatory module in the monocyte-to-macrophage transition that, to our knowledge, has not yet been described in humans. Finally, we show that T2 can also be applied to single cell data, allowing us to identify transcriptional programs in heterogeneous populations, such as asynchronously dividing cells. T2 is a scalable approach to temporal transcriptomics that can be applied to track the activity of thousands of genes in unmodified, primary human cells and tissues, with no genetic engineering.

## Main

The ability to continuously record the fluctuations of the human transcriptome would greatly enhance our understanding of the mechanisms which control it and ultimately, cell fate. RNA sequencing (RNA-seq) has become one of the primary methods for capturing the internal state of cells (*1*). In recent years, tremendous progress has been made in adding a spatial dimension to RNA-seq, allowing us to observe the impact of tissue organization on gene expression. However, much less progress has been made towards adding a temporal dimension: by default, RNA-seq only provides a snapshot of the instantaneous cellular state and fails to capture the dynamics of gene expression over time. In recent years, several technologies have been developed to address this, but all have limitations: molecular recorders order events over long-time frames but require extensive engineering (*2–6*); metabolic labelling measures transcription and degradation rates but requires chemical treatment (*7*); RNA velocity is label-free but provides only the instantaneous derivative of expression in a population of RNA molecules (*8*), rather than a temporal history; and Live-seq enables repeated measurements of living cells but has limited throughput (*9*). There is, to-date, no way to extract detailed information about past transcriptional events at scale from unmodified human cells.

Previously, we described RNA Timestamps (10), which measures the ages of individual engineered RNA molecules by exploiting the accumulation of A-to-I edits. Using a modified adenosine deaminase (ADAR) and an RNA cassette inserted into the 3’UTR of a gene of interest, we measured the ages of single RNA molecules with hour-scale fidelity and reconstructed the past activity of promoters by combining measurements of many mRNAs. However, RNA Timestamps requires extensive genetic engineering and could only measure the activity of a handful of promoters at once, making it impractical for most applications. Here, we show that endogenous A-to-I edits occur throughout the transcriptome in unmodified human cells due to the activity of ubiquitously expressed ADAR enzymes (11, 12), and that these edits are sufficiently widespread to infer transcriptional history across thousands of genes *in vivo* from a single endpoint measurement, with no genetic engineering. By combining the age estimates of individual molecules, we can reconstruct past cellular transcriptional activity in individual cells.

## Endogenous RNA editing encodes RNA age and distinguishes past events

We initially expected that exogenous editors would be necessary to observe significant amounts of editing in most cell types. Thus, we first developed a calibration protocol to identify adenosines across the transcriptome that show significant, time-dependent A-to-I editing (**Fig. 1A**, “editing site”). To identify editing sites, we transfected HEK293 cells with an ADAR2-based editor plasmid (see Materials and Methods) and halted transcription with actinomycin D (ActD, an inhibitor of RNA polymerase II) and then performed bulk short-read RNA sequencing. We screened several editors in this fashion and, with the final editor plasmid we selected (ADAR2(E488Q)-Nlamdba), we identified 428,418 adenosines across 16,168 genes that had a significant increase in editing 8 hours after adding actinomycin D (**Fig. 1B**).

**Figure 1:**
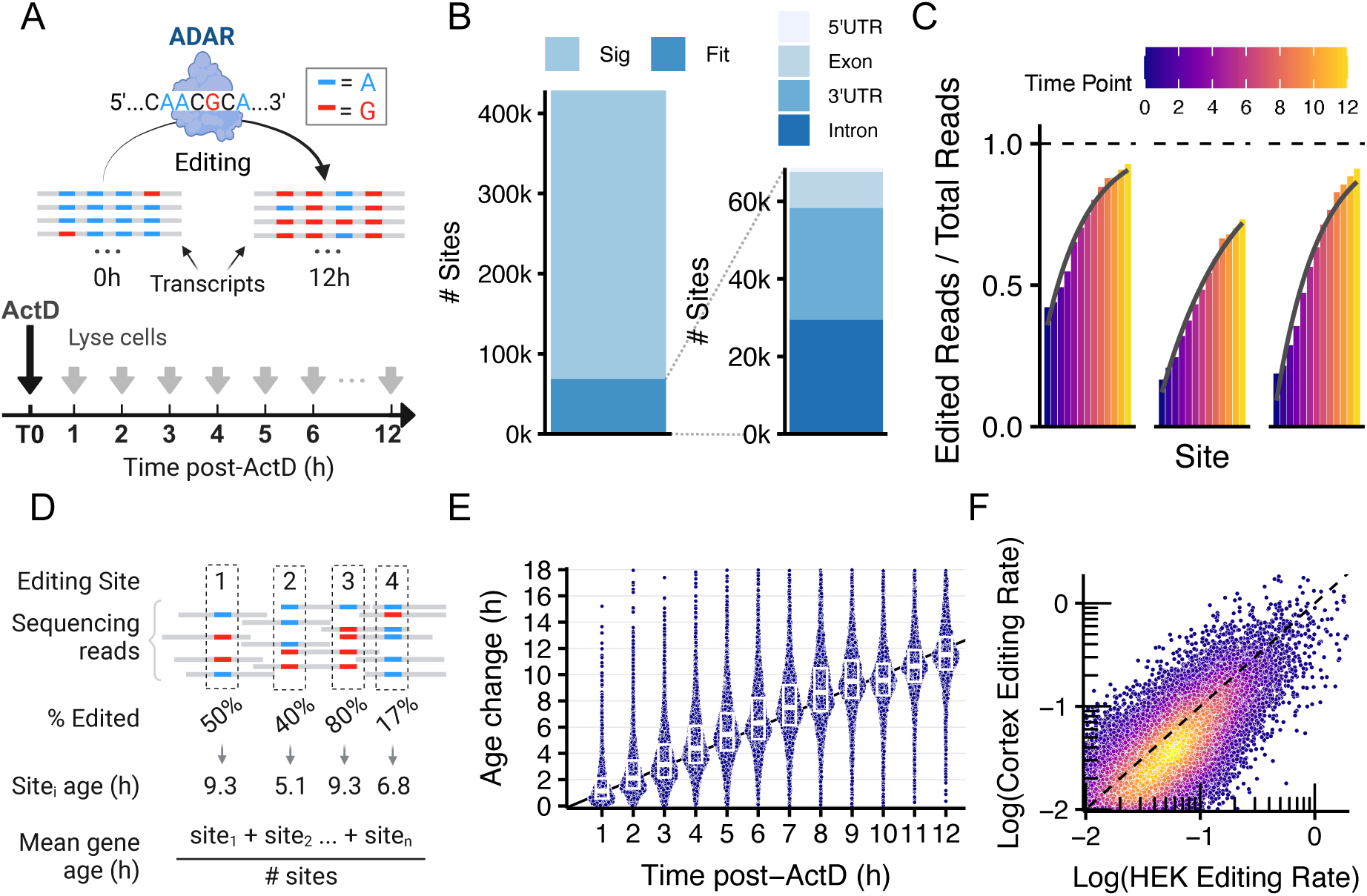
Endogenous editing encodes the age of human transcripts. (**A**) ADAR-catalysed A-to-I edits accumulate over time and the editing rate can be determined by transcription shut off and time-series RNA-seq. (**B**) Left: 428,148 adenosines are significantly differentially edited in HEK cells expressing an ADAR2(E488Q)-Nlambda plasmid 8 hours after adding actinomycin D. Right: Of those editing sites, 68,599 are well fit by an exponential CDF, and their location within genes are shown as stacked bars. (**C**) Editing fractions at three sites in the 3’UTR of the GATC gene are shown at monotonically increasing time intervals following ActD treatment. The fit of an exponential CDF is shown in grey. (**D**) The mean age of a gene’s transcript population can be determined from the editing at each of its sites. (**E**) The mean age of each gene is calculated from the short-read HEK calibration data and the time change compared to the T0 time point (plotted as purple points). A black line of gradient 1 is shown as a guide. Boxplots show the 25th, 50th and 75th quantiles, whiskers extend to the value at most 1.5 * IQR from the respective hinge. (**F**) The correlation of editing rates at shared sites in the ADAR2(E488Q)-Nlambda expressing HEK293 calibration and hiHPSC-derived cortical neuron calibrations are shown as points coloured by 2D kernel density and plotted on log10-log10 axes.

In our previous work (*10*), we found that editing sites were suitable for inferring the age of RNAs if the cumulative editing at those sites was well-fit by an exponential cumulative distribution function (CDF), corresponding to sites that are edited independently and with a roughly constant rate with time since transcription. We refer to this exponential CDF as the editing rate model (**Eq. 1**),

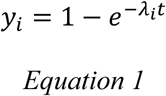

where for site *i, y_i_* is the fraction of edited reads out of total reads (“editing fraction”), *λ_i_* is the editing rate parameter and *t* is the age. We thus collected several timepoints and fit the model in Eq. 1 to all editing sites and found 68,599 editing sites across 6,386 genes were well-fit (R^2^ > 0.4) by an exponential CDF, and that had an editing rate of at least 0.01 per hour (**Fig. 1B,C**, “fit sites”), which is necessary to be useful for downstream analysis. These fit sites had a mean editing rate of 0.058 per hour - corresponding to an expected editing time of 17.2 hours.

Much to our surprise, in subsequent calibration experiments performed in HEK cells with no editing plasmid and in iPSC-derived human neuron cultures (**Fig. S2, methods**), we observed 201,233 fit sites across 8,685 genes. We determined that the effect of the editor plasmid was primarily to dramatically increase the number of editing sites in exonic regions with editing rates that are, in general, too low to be useful for the method (**Fig. S3)**. Thus, in all subsequent experiments, we do not supply any exogenous editor and rely solely on endogenous editing (see Materials and Methods).

We next reasoned that it would be possible to infer the average age of a population of RNAs by inverting Eq. 1, to infer the age *t* from the average editing rate *y* (**Fig. 1D**). We first calculated the average age associated with each fit site, and then combined the average ages for all fit sites across a gene to obtain an age estimate for that gene. Comparing our age estimates to the ground truth by linear regression yielded a slope of 0.98 and a mean coefficient of determination (R^2^) of 0.75 (**Fig. 1E**), suggesting that our endogenous editing loci can faithfully record the average ages of populations of RNA. Moreover, we found the editing rates to be highly correlated between divergent cell types, such as HEK293 cells and hiPSC-derived cortical neurons (Pearson correlation = 0.69, **Fig. 1f**), and could also be detected in human intestinal organoids and primary human brain slices collected from the temporal lobes of patients with focal epilepsy (**Fig. S4**). We refer to the resulting method as Transcriptome Timestamping (T2).

Utilizing improvements in the accuracy of nanopore long-read sequencing (*13–15*), we asked whether it would be possible to infer the ages of individual RNA transcripts by observing the editing states of multiple fit sites on a single molecule (**Fig. 2A**). Conceptually, RNA molecules with many edits are likely to be older while RNA molecules with fewer edits are likely to be younger; in practice, since fit sites for a given gene are sparse in sequence space, we require long-read sequencing to capture multiple fit sites onto a single read. We constructed a likelihood model for the age of a transcript (“single-molecule age”) by modelling each fit site as a sample from a time-dependent binomial distribution (**Fig. 2A**, Methods). We then visualize the “most likely transcriptional history” of a population of transcripts as a histogram of the maximum likelihood estimates (MLE) of the ages of each transcript in the population (**Fig. 2C**). Applying this method to the data from the calibration protocol recovers an increase in the mean ages of genes that accurately tracks the ground truth (slope = 0.86, R^2^=0.61, **Fig. 2D**).

**Figure 2.**
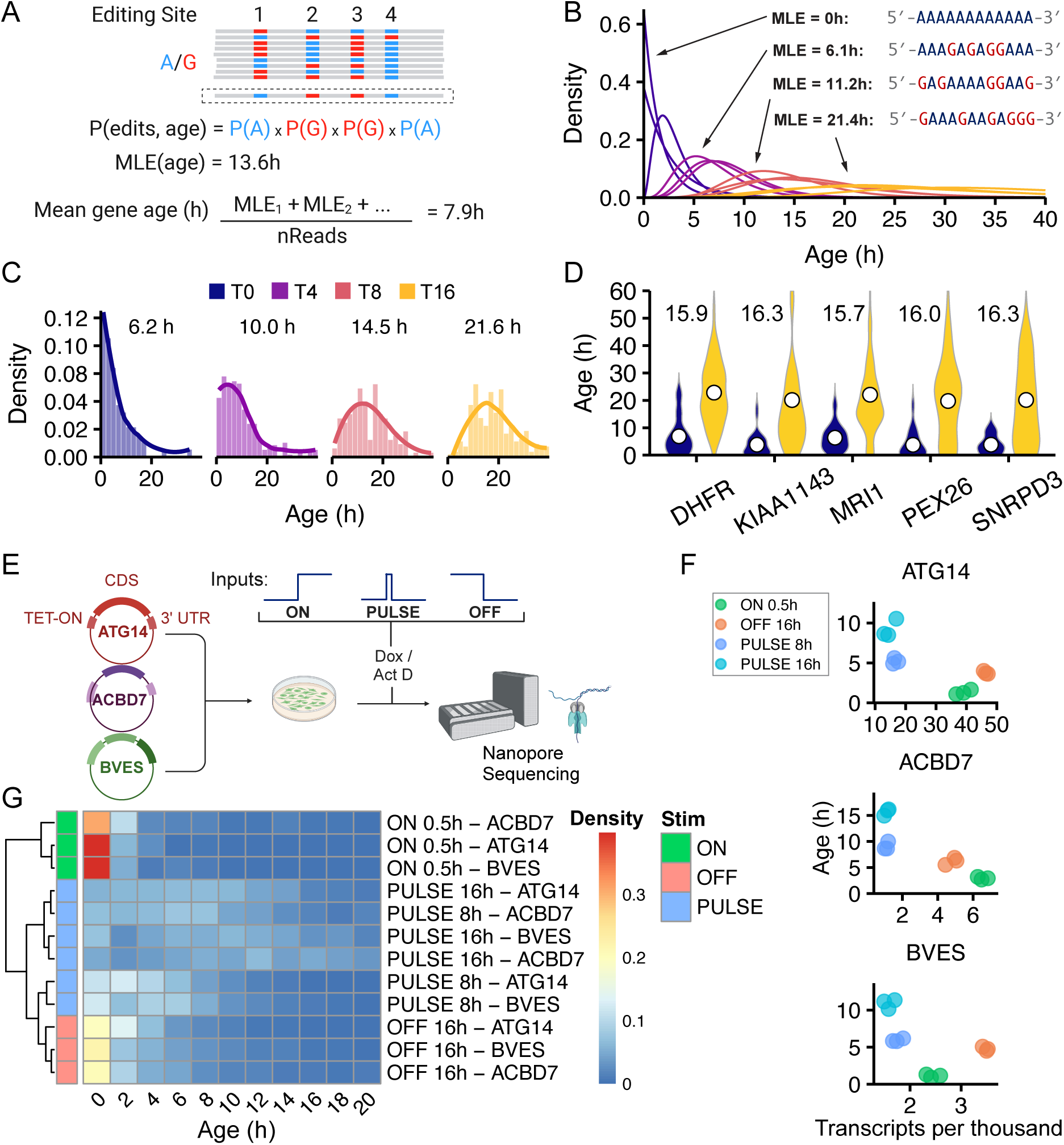
The age of individual endogenous transcripts records transcriptional history. (**A**) The age of an individual RNA molecule can be modelled from the editing state of its sites and summarized by a maximum likelihood estimate (MLE). (**B**) Likelihood functions for the age of individual GATC transcripts from the T0, T4, T8 and T16 time points of the HEK calibration are shown as curves, coloured by time point. MLEs of age for specific reads are given as well as a portion of the editing state. **(C)** The MLEs are calculated for all GATC transcripts and presented as histograms with a non-parametric (LOESS method) smoothed line overlaid. **(D)** MLE distributions are shown as violins for the T0 (navy) and T16 (yellow) calibration time points. The mean age is shown as white circles with the Δ age written above. (**E**) Schematic showing the induction experimental set-up; HEK293 cells were transfected with plasmids containing the genes ATG14, BVES and ACBD7 under the control of the TET-ON system. Three transcriptional programmes were induced using a combination of Dox and ActD: ON, OFF, PULSE 8H and PULSE 16H. (**F**) Points show the mean age and transcripts per thousand values for each of the four behaviours (3 replicates per behaviour). The 3 subplots are for the 3 induced genes. (**G**) Hierarchically clustered heatmap showing the density of the age histograms for each of the conditions for each of the genes. Densities are averaged over the three replicates and histograms are of bin width 2 hours.

We next asked whether T2 can distinguish transcriptional programs from endpoint measurements. To investigate this, we induced four distinct transcriptional programs *in vitro* (**Fig. 2E**). We cloned full-length cDNA sequences of three genes (*ATG14*, *BVES* and *ACBD7*) into TET-ON expression plasmids, transfected HEK293 cells with an equimolar mixture of the three plasmids and then, using a combination of Doxycycline (Dox) and ActD, stimulated them to produce either “ON”, “OFF,” “PULSE 8H,” or “PULSE 16H” behaviors (Methods) before preparing for long-read sequencing. The combination of mean age and mean expression allowed all four conditions to be clearly differentiated from each other, whereas mean expression alone would be unable to distinguish the two “PULSE” conditions, thus demonstrating the utility of age information in differentiating between transcriptional programs (**Fig. 2F**). Moreover, hierarchical clustering on the maximum likelihood age histograms clearly differentiated the four transcriptional programs, with histograms clustering first by stimulation type and then by gene (**Fig. 2G**). Together, these analyses demonstrate the ability of T2 to distinguish complex transcriptional events, either independently or in combination with traditional expression data.

## T2 uncovers transcriptional dynamics in unmodified primary human monocytes

Across Eukarya, only a small fraction of transcription factor (TF)-bound genomic targets identified by CHIP-seq can be validated in perturbation experiments (*16*). One possible explanation for this is that many TFs may have only a transient, facilitatory role in transcriptional regulation. Because T2 relies only on endogenous editing from ADAR enzymes, it may be able to infer transient transcriptional dynamics in unmodified human cells that are not apparent from endpoint RNA sequencing alone, for example because they have no overall effect on transcription levels. To explore this idea, we studied the temporal dynamics of differentiation in human monocytes.

Primary human monocytes freshly isolated from human whole blood were stimulated with E. coli lipopolysaccharide (LPS) and lysed after time intervals ranging from 0 to 6 hours (Fig. 3A). Differentially expressed genes between 0H and 6H were enriched for NFκB, TNF-α and NOD-like signaling pathway members, consistent with differentiation to an ‘M1’ macrophage phenotype (**Table S1**, **Fig. S5**). To identify genes that may be involved in transient transcriptional programs, we developed a new metric that we term ‘differential age,’ which is the age equivalent of differential expression, implemented as the log2 fold change of the mean age between two conditions (Methods). We calculated the differential age for all genes present in the data between the T6 and T0 (baseline) time points in the monocyte differentiation and found 10 genes that were differentially aged but not differentially expressed (**Fig. 3B, Table S2**). To our knowledge, none of these genes were known to have specific roles in monocyte to macrophage differentiation (*17*). This set included, as examples, two ribosomal proteins (*RPS19* and *RPL37A*) that appeared to be significantly older at the 6-hour timepoint despite having no significant change in expression levels. Collection and examination of intermediate timepoints (2h and 4h after stimulation) revealed that, indeed, these genes underwent a transient initial burst of transcription followed by transcriptional shutoff, which was identifiable as an increase in overall gene age despite a return to original levels of expression (**Fig. 3C, S6**). Similarly, we identified a cytochrome p450 orphan gene (*CYP20A1*) that had no significant change in abundance at any of the time points (**Fig. 3D**), but that nonetheless had a significant decrease in age at the 6-hour timepoint. We hypothesize that the shift towards younger transcripts may be caused by a concomitant increase in both transcription and degradation rate. In support of this, *CYP20A1* transcripts are predicted to harbour at least 10 miRNA recognition elements, and miRNA overexpression experiments suggest that *CYP20A1* transcript stability is indeed regulated by miRNAs (*18*, *19*). Additionally, monocytes treated with LPS for 6 hours are known to upregulate 7 miRNAs which are predicted to bind to the 3’ UTR of *CYP20A1* (*20*) (see Table S3). Thus, T2 enables the identification of transient gene expression programs that are not identifiable from endpoint expression measurements alone.

**Figure 3.**
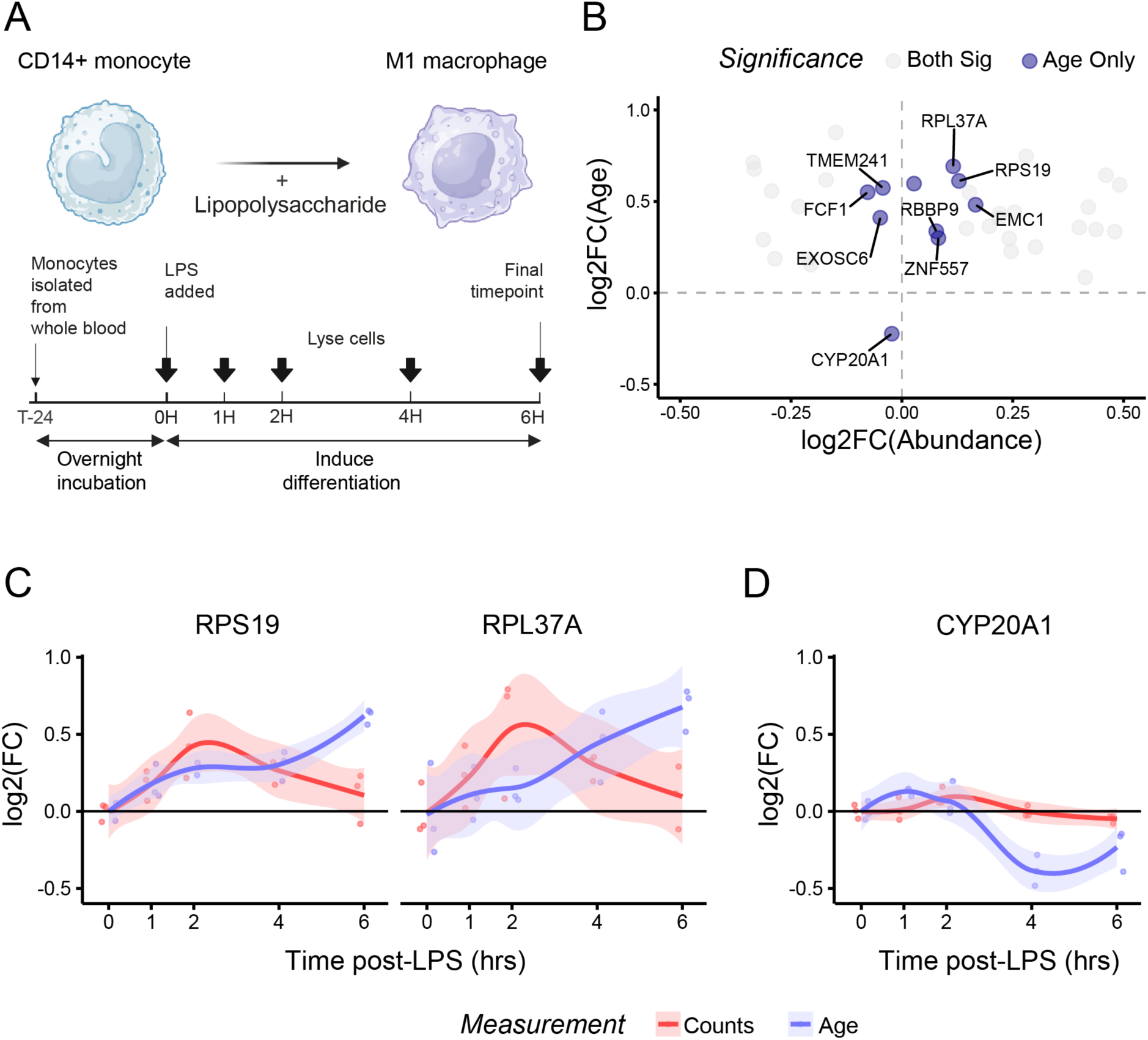
T2 records transcriptional history in primary human monocytes. (**A**) Schematic of the isolation of CD14+ human monocytes, stimulation with 18/11/2024 21:14:00LPS and time-series RNA-seq. (**B**) Log2 fold changes in age (y axis) and abundance (x axis) for genes that are either significantly different in age only (purple) or both age and abundance (grey) 6 hours post-LPS stimulation. Age only genes are labelled. (**C**) RPS19 and RPL37A have transient up/down-regulation at intermediate timepoints, (**D**) CYP20A1 has a change in age despite no significant change in abundance at any time point.

## T2 identifies transcriptional modules and gene regulatory pathways

We next asked whether T2 could be used to identify new transcriptional modules (i.e., genes with similar temporal dynamics) in monocyte differentiation. Our approach, which relied on hierarchical clustering with a bootstrapped robustness threshold on the differential age data (Methods), revealed ten clusters (**Table S3**), of which we identified four for subsequent analysis. Genes in cluster 1 undergo a modest increase in age after LPS induction before stabilising (**Fig. 4A**). Genes in cluster 2 undergo a transient decrease in mean age, followed by an increase, possibly indicative of an immediate-early program. Finally, clusters 3 and 4 appear to undergo gradual and continuous increases and decreases in mean age, respectively.

**Figure 4.**
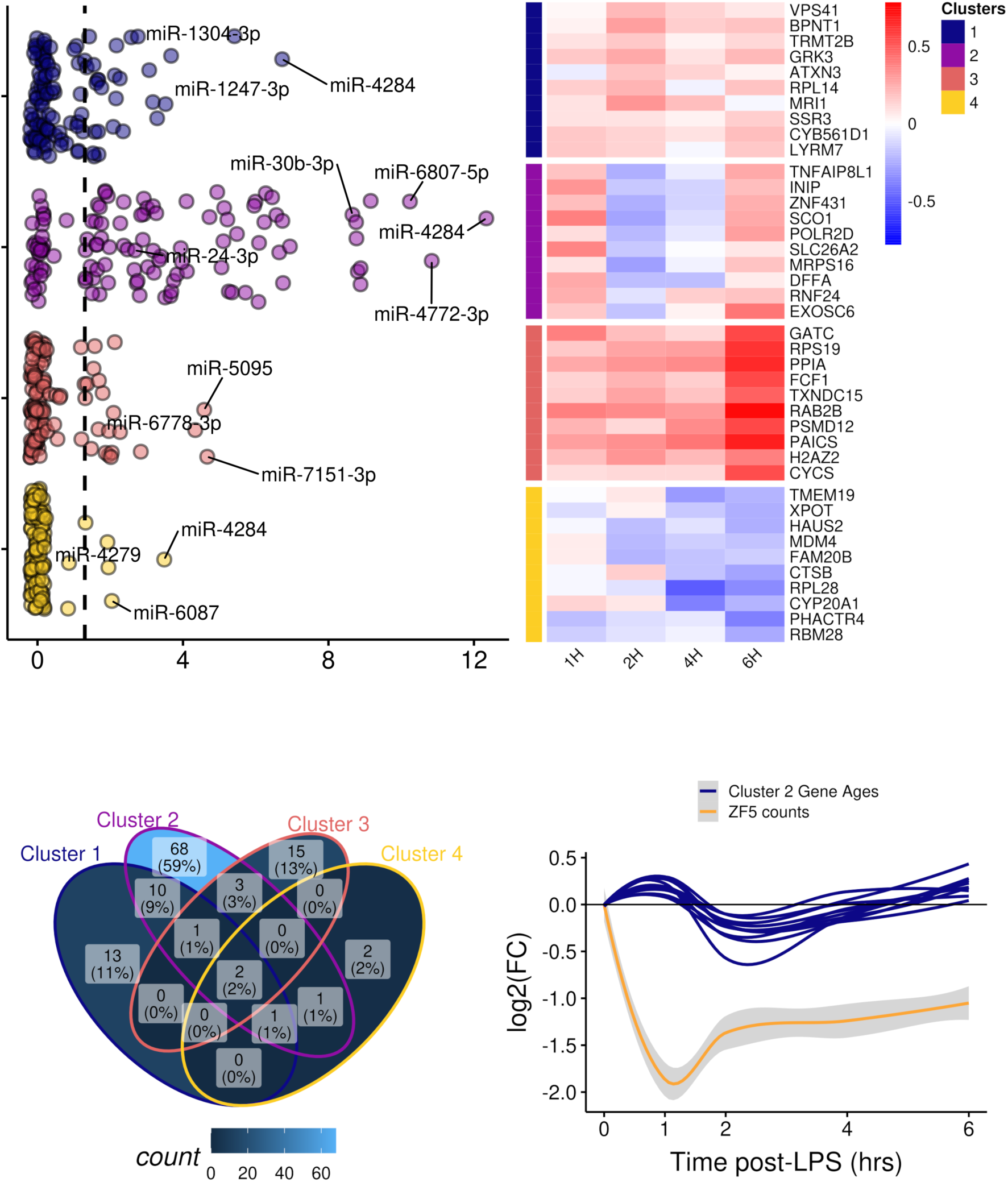
Differential age identifies clusters of genes enriched for different sets of miRNAs and TFs. (**A)** Differential Age: Genes were clustered by differential age relative to the 0H timepoint. Four clusters of interest are shown in a heatmap of differential age. Differential age is shown as log2-fold change in mean age. The 10 genes closest to the centroid of each cluster are shown. miRNAs: Jitter plots display the −log10(p-adj.) values of enriched miRNAs for each cluster. The red dashed line shows significance (p<0.05). Labels are assigned to miRNAs of interest which are discussed in the main text. (**B)** Venn diagram showing the sharing of significantly enriched miRNAs between clusters 1-4. (**C)** For each of the 10 genes displayed for cluster 2, the relationship between the gene’s age and ZF-5 counts for the duration of the LPS stimulation.

Co-aging genes may share common transcriptional regulatory mechanisms such as miRNAs and transcription factors. To investigate this, we applied transcription factor and miRNA enrichment analysis to the clusters. Interestingly, we found that all 88 genes in cluster 2 were in proximity to a ZF-5 regulatory motif (**Fig. 4B**, **Table S3**). ZF-5 (ZBTB14) is a TF known to regulate the differentiation of monocytes in zebrafish, but whose role in human monocyte differentiation is not known. In zebrafish, ZF-5 represses the transcription of PU.1, a master regulator of monocyte differentiation (*21*), thereby limiting monocyte proliferation, and a loss-of-function mutation in the *ZBTB14* gene (ZBTB14S8F) has been identified in a patient with Acute Myeloid Leukaemia (Tyner et al. 2018), suggesting a similar function in humans. To investigate this further, we examined intermediate timepoints and observed that the expression levels of ZF-5 indeed fall initially before recovering (**Fig. 4C**), mimicking the changes in mean age in cluster 2. This suggests that cluster 2 genes represent part of a transcriptional unit which regulates monocyte differentiation, which themselves are regulated by ZF-5.

A similar analysis also identified numerous microRNAs that are likely to be involved in the monocyte to macrophage differentiation program (Fig 4A and 4B). Our analysis identified 68 miRNAs that were significantly associated with genes in cluster 2. This included several miRNAs that are known to be upregulated in LPS-treated human monocytes, such as miR-4284, the most highly enriched miRNA in cluster 2, and the miR-30b/miR-24 duo, which are both known to temper the production of cytokines and phagocytic capacity in human macrophages (*22*, *23*). We also identify several novel putative miRNA modulators of monocyte differentiation. For example, miR-5095 is the most highly enriched miRNA in cluster 3 and has not previously been associated with human monocyte differentiation. Interestingly, the miR-5095 mouse ortholog belongs to the super-enhancer of genes which determine CD14+ monocyte cell identity, suggesting it may play an undisclosed role in human monocyte-to-macrophage differentiation (*24*). Together, these results demonstrate the ability of T2 to identify new gene regulatory pathways in transient and transitional transcriptional programs.

## T2 detects transcriptional dynamics in single cells

Finally, because T2 functions at a single-molecule level, we asked whether it could be used to interrogate transcriptional histories in single cells. We used SMARTseq2 and a PromethION24 (ONT) to perform single cell long-read sequencing on 96 single HEK293 cells (**Fig. 5A**).

**Figure 5.**
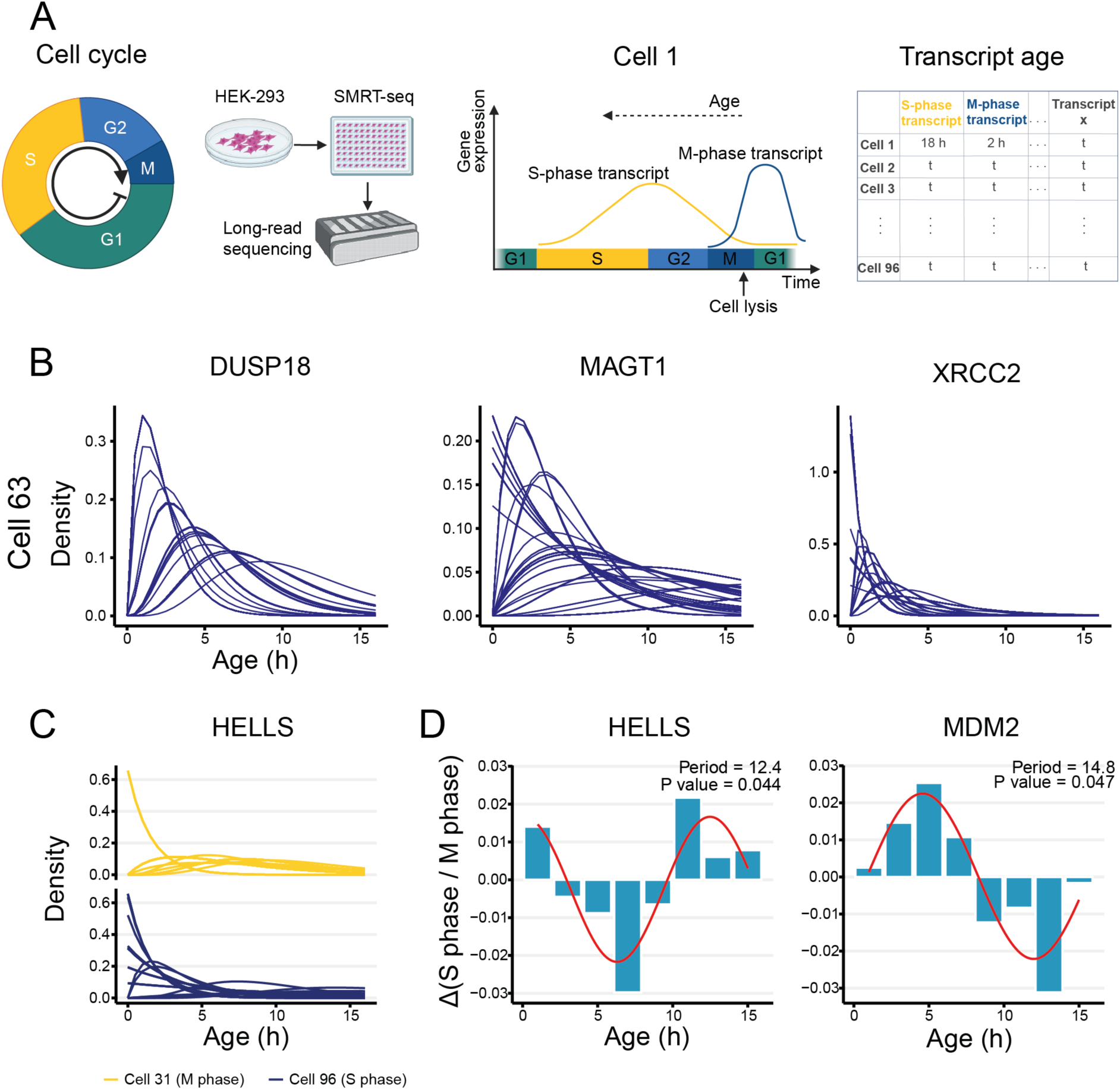
T2 resolves transcript age at single cell resolution. (**A**) Schematic showing experiment design, deriving transcript age at single cell resolution. (**B**) Age distribution of transcripts (individual mRNA molecules) for 3 genes in a single HEK293 cells, *DUSP18*, *MAGT1*, *XRCC2*. (**C**) HELLS transcript age likelihoods for cell 31 (M-phase, top) and cell 96 (S-phase, bottom). (**D**) Delta-density of top 15 S-verse M-phase cells per time-bin for HELLS and MDM2, with fitted sine curve (red). Text shows derived dividing time of cells (Period) and p-value from 750 bootstraps of the residual standard error from non-linear least squares regression.

As expected, T2 resolved age distributions from individual transcripts in single HEK293T cells (**Fig. 5B**), and correctly identified known cell-cycle markers such as *HELLS* as being younger in G1/S-phase cells and older in M-phase cells (**Fig. 5C**). We asked whether T2 could additionally extract the periodicity of the cell cycle from our mixed population of single cells. We used non-linear least squares regression to fit a sine curve to the relative densities of expression of 34 genes with sufficient read depth in S-versus M-phase cells and used a bootstrapping approach to evaluate significance of the resulting fits (Methods). Remarkably, we found *HELLS* to have a statistically significant fit (p=0.044) with a period of 12.4 hours, while *MDM2* was also expressed in a cyclical fashion (p=0.047) with a period of ∼14.8 hours (**Fig. 5D; Table S7**). The phase of MDM2 was offset by about 120 degrees from the phase of HELLS, consistent with known roles of *MDM2* in the cell cycle, both as an inhibitor of p53 at the G1/S transition point (*25*), but also in promoting cell-cycle progression throughout G1 phase via *E2F-1*, *p21* and *Rb* (*26*).

By leveraging the endogenous editing activity of human ADAR1, T2 provides a method to extract complex temporal information from long or short read RNA sequencing across thousands of genes in unmodified human cells. T2 should be applicable to any human tissues or cell types that express ADAR1, which includes most tissues, and in principle can even be applied retroactively to previously gathered datasets. Subsequent work on second generation exogenous editors identified candidates, such as TadA, that could greatly increase the resolution and coverage of T2 by increasing the rate and abundance of RNA editing throughout the cell, without immediately apparent effects on cell physiology (**Fig. S8**). Moreover, several natural extensions of T2 are possible, such as extensions to other post-transcriptional modifications, which may increase time resolution and coverage. We suggest that T2 may be a valuable part of the RNA sequencing toolbox for inferring transcriptional dynamics.

## Acknowledgments

We acknowledge Gabrielle Chappell for clinical sample processing; James Evans and Minee Lee Choi from the Ghandi lab who provided us with hiPSC-derived cortex cultures; Rory Maizels for providing the empty TET-ON plasmid used in Fig. 2 in addition to some of the editing construct plasmids used in Fig. S1 (See Table S4 for details); Danny Lang from the Scientific Computing STP for help with high-performance computing applications; Daniel Snell from The Francis Crick Advanced Sequencing STP for support with Nanopore sequencing; Neelam Mehta, Danson Loi and Eleanor Calcutt from the Cribbs Lab at Oxford University for help with long read sequencing; Molly Strom, Karl Dietrich Brune, Daniele Cervettini and Oscar Wilkins for technical and intellectual support throughout the project; the Long Read Sequencing Facility at UCL for providing access and support with PacBio sequencing; and the following Science Technology Platforms at the Francis Crick Institute: Flow cytometry, Cell services, Advanced Sequencing, Scientific Computing.

## Author Contributions

S.G.R. conceived of the project. A.G., J.B., and A.W. performed the experiments and analysis. A.G., J.B., A.W. and S.G.R. wrote and edited the final manuscript. A.M.A performed cell culture and assisted with single cell experiments. D.M. performed data analysis and visualisation on the calibration datasets. A.P.C. provided reagents, instrumentation, computing resources and advice on Nanopore library preparation. L.M. provided expertise and assistance with gut organoid culture. S.G. provided the hiPSC cell lines. M.R., S.G., E.G., and M.C., provided reagents, support, commentary, and edits. L.C. and G.Y. provided support with methodology and model design. S.G.R. supervised and managed the project and secured funding.

## Data and Materials availability

All plasmids are available upon request. Original code and data structures have been deposited at GitHub and are publicly available (reference link to repo).

## List of Supplementary Materials

Materials and Methods

Supplementary Text Figs. S1 to S8

Tables S1 to S3

References (26–32)

## Supplementary Materials

### Materials and Methods

#### Cell Culture

##### HEK293

HEK293-FT cells were cultured in DMEM (Thermo Fisher, 11965092) supplemented with 10% FBS (Gibco) and 1% Penicillin-streptomycin (Sigma). HEK293T cells were seeded on tissue culture plastic 24-well plates in triplicate 48 hours before the experiment.

##### Human hiPSC derived neurons

Human induced pluripotent stem cells (hiPSCs) were generated from reprogrammed fibroblasts from healthy donors, with approval from the London-Hamstead Research ethics Committee and the University College London, Great Ormond Street Institute of Child Health and Great Ormond Street Hospital Joint Research Office. The hiPSCs were cultured on Geltrex (Thermo Fisher) in E8 media (Thermo Fisher) or mTeSR (Stem Cell technologies) and passed using 0.5 mM ethylenediaminetetraacetic acid (Thermo Fisher). Neurons were generated using an established protocol (*27*). Briefly, neocortical stem cell differentiation was achieved with dual SMAD inhibition using SB431542 (10µm, Tocris) and dorsomorphin dihydrochloride (1µm, Tocris) for 12 days, followed by ongoing culture for an additional 90 days to differentiate into neurons.

##### Human Organotypic Brain Slices

Organotypic brain slice cultures were prepared using the interface method adapted from Ravi et al. (*28*) and De Simoni & Yu (*29*). Fresh healthy cortical human brain tissue was removed during neurosurgery and used for research under appropriate National Research Ethics approval (Reference: 21/SC/0111). The tissue was resected and place in ice-cold dissection media in the operating theatre and transferred for immediate sectioning at 300uM thickness using a Leica VT1200s vibratome. Each 1cm2 brain section was plated on a 30mm culture plate insert (Millipore, PICM03050) in a 35mm Nunc 6-well plate flooded with 1ml of culture media per well, and transferred to an incubator at 37 °C and 5% CO2. Media was changed every 24 hours. Dissection media was composed of Hank’s balanced salt solution (HBSS) supplemented with HEPES (pH 7.4, 2.5 mM), D-glucose (30 mM), CaCl2 (1 mM), MgCl2 (1 mM), and NaHCO3 (4 mM). Culture media was composed of Neurobasal l-Glutamine supplemented with 2% serum-free B-27, 2% Anti-Anti, 13 mM d-glucose, 1 mM MgSO4, 15 mM HEPES, and 2 mM GlutaMAX. RNA extractions were performed in triplicate using the Qiagen RNeasy Plus Universal Mini Kit, following manufacturer protocols.

##### Organoid Culture

Cultures of human intestinal organoids were established as previously described (*30*). Briefly, organoids were re-established in culture and maintained as three-dimensional spheroids in CultrexTM reduced growth factor basement membrane extract, type 2 (R&D Systems) from previously cryopreserved organoids originally derived from human intestinal biopsies (*31*) (Research Ethics Committee references 04-Q0508-79 and 18/EE/0150). Organoids were cultured in 24 well plates using human IntestiCult Organoid Growth Medium (STEMCELL Technologies) supplemented with 3μM CHIR99021 during expansion phase, with a passaging ratio of 1:5 and media changes every 2 days. For hypoxia experiments, organoids were treated with 1mM of Dimethyloxallyl Glycine (DMOG) for 6 hours. For necroptosis experiments, organoids were stimulated with 30ng/ml of recombinant TNF-alpha (R&D Systems) for 6 hours. RNA extractions were performed in triplicate using the Qiagen RNeasy Plus Mini Kit, following manufacturer protocols.

#### Calibration Protocol

Cells (either HEK293s or hiPSCs) were cultured as previously detailed. Actinomycin D was added to the cells to a final concentration of 1ug/ml in complete media at time t=0 hours. Wells were lysed in triplicate at timed intervals. For example, the HEK293 cells were lysed at the following timepoints: (0,1, 2, 3, 4, 5, 6, 7, 8, 9, 10, 11, 12, 16, 24, 38H).

RNA was extracted using the RNEasy Plus Mini Kit (Qiagen 74136) as per the manufacturer’s instructions. Cells were lysed in the wells using RLT plus from the Qiagen kit and vortexed to homogenise. Purified total RNA was then quantified using the Qubit fluorometer (Life Technologies Q32855) and quality was measured on an Agilent Tapestation using RNA Screentape (Agilent 5067-5576).

Full-length, short-read sequencing libraries for Illumina were then prepared by poly-A selection using oligo-dT magnetic beads (NEB E7490) followed by the Ultra II directional library prep kit for Illumina (NEB E7760) according to the manufacturer’s instructions. In brief, RNA is fragmented, a library is prepared by reverse transcription with random primers and adapters are ligated to both ends of the cDNA. Uracil is incorporated during second strand synthesis and strand specificity is achieved by digestion of the second strand by USER. Sequencing adapters and sample barcodes are added by PCR before purification, QC, and pooling. Short read sequencing was performed on a NovaSeq (Illumina) using 150 cycle kits with 76 bps for read 1 and 2. The samples were sequenced at a depth of approximately 225 million reads per timepoint (HEK293) and 500 million reads per timepoint (Cortex).

Long-read sequencing of the HEK293 and cortex samples was performed using the PacBio Iso-seq on a Sequel IIe. Library prep used the Iso-Seq Express Template Preparation kit (PacBio) according to the manufacturer’s instructions. Both timepoints in the Cortex calibration were sequenced on one Sequel IIe SMRT Cell 8M flow cell in CCS mode at a mean depth of approximately 2.2M reads. In subsequent long read sequencing experiments, the Oxford Nanopore Technologies platform was utilised given its higher throughput, with similar performance in determining single-molecule ages (below).

#### Human gene induction experiment

##### Cloning

Total RNA was extracted from HEK293 cells using the RNEasy mini kit and a cDNA library was generated. ATG14, BVES and ACBD7 were then amplified from this cDNA library using transcript-specific primers. Bands were then gel-purified and assembled into a plasmid vector using NEBuilder Hi-Fi DNA assembly (E2621). E.cloni 10G Chemically Competent Cells (Lucigen) were transformed and grown overnight. The sequencing data allowed us to identify pre-edited ADAR editing sites. PCR was used to restore these editing sites to their unedited form. The PCR products from these reactions were then reassembled using NEBuilder Hi-Fi assembly and validated using whole plasmid sequencing. The final products were plasmids, each containing the full cDNAs of the genes ATG14, BVES and ACBD7, respectively. These plasmids are known henceforth as pAG051, pAG053 and pAG054, respectively.

##### Cell transfection

HEK293 cells were transfected using the Transit-X2 delivery system (Mirus Bio) when the cells reached 70% confluency. Each well of a 24-well plate received 250ng of tTA3 transactivator plasmid and 250ng of an equimolar mixture of the three mammalian cDNA plasmids (pAG051, pAG053 and pAG054). Transfection was performed 48 hours before the t=0 timepoint. The ON, OFF and PULSE conditions were performed as follows. **ON**: at t=0 hours, doxycycline was added. Wells were lysed at 0, 0.5, 1, 4 and 8 hours. A mock induction was lysed at 8 hours. **OFF**: Doxycycline was added at 24 hours before the t=0 timepoint. Cells were changed to doxycycline-free media at t=0 hours and then lysed at t=16 hours. **PULSE**: at t=0 hours, doxycycline was added. ActD was added at t=4 hours. At t=8 and 16 hours, the cells were lysed. For each of these conditions, doxycycline was added to a concentration of 1ug/ml. Wells were lysed in triplicate at each timepoint.

##### Library preparation and sequencing

Library preparation was undertaken using a bespoke SMARTseq protocol designed to minimise the number of PCR cycles used to minimise PCR errors (courtesy of Adam Cribbs and Danson Loi). In brief, at least 100ng of total RNA was denatured at 65°C in the presence of 100uM poly-T reverse primer and 1mM dNTPs. Reverse transcription was performed by adding RT buffer (Thermo Fisher), Maxima H-reverse transcriptase (Thermo Fisher) and template-switching oligonucleotide (Table S5). A bespoke temperature ramp-up and cycle programme was used (Table S6). Excess primers were digested by incubation with heat labile Exonuclease I (Thermo Fisher). Sample clean-up was performed with SPRI-select beads (0.6X by volume). Initial PCR was performed with 2X KAPA ReadyMix (Roche) and ONT PCR handle primer. This was followed by a second PCR with the reaction split into 4 (Table S6).

Samples were prepared for sequencing using the ligation sequencing kit v.14 (ONT, SQK-LSK-114) as per the manufacturer’s instructions. Samples were sequenced on a Promethion24 sequencer (ONT) over 5 R10.4.1 flow cells yielding an average of approximately 70Gb per flow cell.

#### Monocyte LPS stimulation experiment

##### Peripheral Blood Mononuclear cell (PBMC) isolation

Whole blood was sourced from a healthy male via the UK National Health Service blood bank. Blood volume was standardized by supplementing with Hank’s Balanced Salt Solution (HBSS) containing 0.3mM EDTA to a final volume of 25ml. Next, 20ml of Ficoll-Paque was dispensed into separate 50ml tubes. The blood was delicately layered atop the Ficoll using a 5ml pipette, ensuring the preservation of the discrete interface. The sample was centrifuged at 700g, without brake, for 25 minutes at room temperature. Post-centrifugation, the PBMCs, identifiable as a white interphase layer, were harvested using a Pasteur pipette and transferred into a clean 50ml tube. The cell suspension was diluted to 50ml with HBSS 0.3mM EDTA, followed by a centrifugation step at 500g for 10 minutes at room temperature. The supernatant was decanted, and the cell pellet was subsequently resuspended in 40ml HBSS 0.3mM EDTA. This wash step was repeated once. After the final centrifugation, the PBMCs were resuspended in 1ml of MACS buffer in preparation for CD14+ isolation.

##### CD14+ cell isolation

Cells were processed using the MagniSort system (Invitrogen). Initially, 200µl of MagniSort Enrichment Antibody was added and thoroughly vortexed. The mixture was incubated at room temperature (RT) for 10 minutes. Subsequently, 3ml of MACS buffer was introduced, followed by centrifugation at 300g for 3 minutes at room temperature. The pellet was resuspended in 1ml MACS buffer. Next, 300µl of MagniSort Positive Selection Beads was added and the solution was vortexed. Following another 10-minute incubation at RT, the volume was adjusted to 2.5ml with MACS buffer and mixed by pipetting. The tube was placed in a magnet and incubated at room temperature for 5 minutes. The supernatant, containing CD14-cells, was discarded and the magnetic separation repeated twice more after resuspending the beads in MACS buffer. The retained CD14+ cells were supplemented with 30ml Iscove’s Modified Dulbecco’s Medium containing 10% Fetal Bovine Serum, 1% Penicillin-Streptomycin-Glutamine and uM GM-CSF.

##### LPS induction

Human CD14+ monocytes were plated in triplicate in 6-well plates at a density of 2 million cells per well immediately after magnetic isolation and 24 hours before LPS stimulation. At time t=0 hours, E. coli (LPS) purified by ion-exchange chromatography (Sigma L3024) was added to each experimental well to a final concentration of 100ng/ml.

##### Library preparation and sequencing

Samples were reverse transcribed and amplified using the same protocol as with the HEK293 induction experiment (see above). They were sequenced on a Promethion24 machine (ONT) over eighteen R10.4.1 flow cells (one per sample) at an average depth of approximately 70M reads each - yielding over 1.2Bn long reads.

#### Single cell RNA sequencing in plates

HEK293FT cells (Thermo Fisher) were grown to 70% confluency, trypsinised, counted and resuspended in FACS buffer (PBS, 2% molecular grade BSA, 1ug/ml actinomycin D) to a final concentration of 5E6 cells per ml. Shortly before cell sorting, DAPI was added as a viability dye. Before the first sort, the FACS machine was calibrated using a horseradish peroxidase assay to ensure that cells were deposited into each well. Single cells were sorted into single wells of a low bind 96-well plate (Eppendorf) using the MoFlo XDP cell sorter (Beckman Coulter) using a 100um nozzle and the following lasers and filters: 405nm, 447/60. After gating on FSC/SSC and a doublet gate, DAPI negative cells were selected for sorting.

The NEBNext Single Cell/Low Input cDNA Synthesis & Amplification Module (NEB E6421L) was used to prepare SMARTseq libraries. Cells were sorted into wells containing 5ul of fresh lysis buffer. Immediately after sorting, samples were prepared for reverse transcription and library preparation. After reverse transcription and 22 cycles of PCR amplification as per the manufacturer’s instructions, an additional 5 cycles of PCR amplification were undertaken to add nanopore barcodes and adaptors. Samples were then pooled and prepared for sequencing using the ligation sequencing kit V14 (ONT, SQK-LSK-114) as per manufacturer’s instructions.

The library was sequenced on a Promethion24 machine (ONT) over six R10.4.1 flow cells with a mean output of 44M reads per flow cell

#### Hyperactive editor experiments

HEK293 cells were transfected with the editors (and corresponding guide RNAs if required). Twenty-four hours later, we added actinomycin-D to stop transcription and lysed cells at 0-hour and 8-hour timepoints in triplicate. In the first generation of editor selection, we studied the full-length hyperactive ADAR2 mutant ADAR2(E488Q). Additionally, several fusion proteins of the ADAR2(E488Q) catalytic domain were tested: ADAR2(E488Q)-Nlambda (*32*), SNAP-ADAR2(E488Q) (*33*), Cas13-ADAR2(E488Q) (*34*), and PABP-ADAR2(E488Q) (courtesy of Rory Maizels). We also tested ABEmax (*35*) and the cytosine base editor BE3 (*36*). Although SNAP-ADAR2(E488Q) showed slightly higher levels of editing than ADAR2(E488Q)-Nlambda, we decided not to proceed with it because its guide RNA cannot be genetically encoded due to the need for an O6-benzylguanine base.

For the second generation of editors, we used a chassis based on the E. coli tRNA-specific adenosine deaminase (TadA). Plasmids for ADAR2(E488Q)-Nlambda (the best construct from generation 1), TadA7.10 (*37*), TadA8.20 (*38*), TadA8.20–Nlamdba and TadA8.20–Dps-N (*39*, *40*) (Dps-N construct courtesy of Karl Brune) were transfected into HEK293 cells and the experiment performed as for the first generation editors.

#### Bioinformatics

Unless otherwise specified all analysis was undertaken in R4.1.0 (https://www.r-project.org/).

##### Short-read data processing

Bcl files resulting from short-read sequencing (Illumina technologies) were converted to FASTQ files by blc2fastq2 v2.20.0 (https://emea.support.illumina.com/downloads/bcl2fastq-conversion-software-v2-20.html). The fastq files were trimmed with cutadapt v3.5 (https://github.com/marcelm/cutadapt/) and aligned to the GRCh38.100 human reference genome using HISAT2 v2.1.0 (http://daehwankimlab.github.io/hisat2/). The counts per gene were then quantified using salmon v1.4.0 (https://combine-lab.github.io/salmon/). Editing was quantified using JACUSA2 v2.0.4 (https://github.com/dieterich-lab/JACUSA2) with call-1 (detect) with settings to exclude any potential editing sites near the start and end of reads, indel positions and splice sites, as well as sites within homopolymer runs of more than 7 bases. The resulting JACUSA files were converted to BED files using a custom python3 code.

##### Long-read data processing

POD5 files from ONT sequencing were basecalled in Super Accuracy Mode (SUP) using guppy_basecaller from guppy v6.4.6-CUDA-11.7.0 on NVIDIA A100 and V100 graphic processing units. The resulting FASTQ files were demultiplexed using guppy_barcoder from the same version.

Demultiplexed fastqs from ONT or PacBio were subsequently run through the same pipeline. Reads were aligned to the GRCh38.100 human reference genome using minimap2 (https://github.com/lh3/minimap2) and quantified using IsoQuant v3.0.0 (https://github.com/ablab/IsoQuant). Edits were quantified as for short-read data (above).

For single-molecule age estimation, base calls at specific loci were pulled directly from .bam files using custom R code that is made available in the github repo of this project in the script called extract_calls.R.

Long-read data from single cells was deduplicated using samtools v1.13 markdup (https://www.htslib.org).

#### Modelling and Statistics

##### Editing rate determination from calibration data

Sites with significant increases in editing 8 hours post-ActD addition were found using JACUSA2 call-2 and filtering to sites with Z score > 1.96 (‘Significant sites’). A linear model was then fit to each site and any sites that had a non-positive gradient were removed (indicating either no accumulation of edits or a potential SNV).

Since the fraction of reads that are edited at each site is likely non-zero at the 0h calibration time point, the editing rate equation:

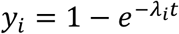

was modified with an additional parameter, *a*, to yield the calibration equation:

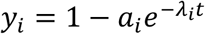

where for site *i*, *y* is the fraction of reads that are edited, *a* is a free parameter that approximates the complement of the fraction of edited reads at the start of the calibration experiment, *λ* is the editing rate and *t* is the time since the start of the calibration experiment.

The calibration equation was then fit to each of the remaining sites by non-linear least squares using the nls function from the R package *stats* v4.1.2 with starting values *λ*=0.01 and *a* = 1 − *y*(*t* = 0). The resulting list of editing sites were then filtered on the following criteria to give the list of ‘fit sites’:

- R^2^ ≥ 0.4
- 2 ≥ *λ* ≥ 0.01
- Each site must appear in at least 4 time points from the calibration dataset.
- The predicted age of the site at the T0 calibration time point is less than 25 hours.

Sites were annotated as belonging to a specific gene if their genomic coordinates mapped between the start of the 5’UTR and the end of the 3’UTR and matched the sense of the strand. Sites that were present in regions that are annotated with more than one gene on the same strand could not be assigned unambiguously and were removed from analysis. Sites that mapped to intergenic space were also removed from the list of fit sites.

The calibration datasets from hiPSC-derived cortical neurons and HEK293 cells (transfected with ADAR2(E488Q)-Nlambda) were compared by taking the logarithm base 10 of the editing rates and calculating the Pearson correlation coefficient for the shared sites. (Since we found the editing levels in HEK293 cells transfected with ADAR2-Nlambda and un-transfected HEK293 cells to be similar, we used the ADAR2-Nlambda calibration data for all experiments in this manuscript as it was performed on more timepoints than the un-transfected HEK293 experiment.) The datasets were combined by calculating the mean log fold change at all shared sites, scaling the editing rates at the hiPSC-only sites by this factor and then appending them to the list of HEK293 editing sites. The total set of 201,233 fit sites used for analysis was obtained from this combined dataset as described above.

##### Per-site age

Given a pre-determined editing rate (*λ*) and an observed fraction of edited reads (*y*), the editing rate equation was rearranged to find the mean age (*t*) of the population of transcripts that contain a particular site *i*:

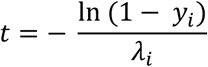

To obtain the mean age of a given gene, the mean of the per-site age estimates over all the sites annotated to that gene is taken.

To investigate how the mean age changes over the course of the calibration experiment, for each gene (n = 5,540) we took the mean of the per-site ages at all time points and subtracted the age at T0. If the gene had no sites in the T0 timepoint then it was removed. To avoid infinitely old sites, any sites that were edited on all reads were modified by subtracting 1 / the number of reads and the age then taken.

##### Single-molecule age

From long-read sequencing .bam files we extract the base calls at fit sites and model the age of individual transcripts by constructing a likelihood function for the age of the transcript given the editing state, *x*, over its sites

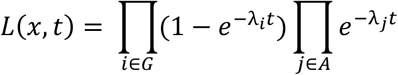

where set *G* contains the sites that are edited and set *A* contains the sites that are not edited.

When calculating single molecule age, we kept only those genes that had at least 5 fit sites in the 3’ UTR, resulting in 32,803 sites across 949 genes. We obtained point-estimates for the age of the reads via maximum likelihood estimation (MLE) performed with the maxLik function (R package *maxLik* v1.5-2) on the log of the likelihood function for each transcript.

We estimated the population distribution for any given gene by calculating a histogram of the MLE values (typically between 0 and 20 hours with a bin width of 2 hours). Depending on the analysis, we calculated either a histogram with raw counts of MLEs per bin or a density where each bin is normalised by the total counts. When shown in conjunction with a smoothed line, the line was generated using *ggplot2* v3.4.1. with the LOESS (locally estimated scatterplot smoothing) method. When shown as connected points with error bars, the points show the mean of the bin values across the replicates and the error bars denote the standard error of the mean of the replicates.

##### Differential age

We calculated the differential age between two conditions by taking the log2FC of the mean age between the conditions and determined the significance using a Mann-Whitney U test on the MLE values (replicates are pooled prior to testing). For each contrast tested, p values were adjusted using the Benjamini-Hochberg method.

##### HEK293 Tet-on induction analysis

For each condition the age was calculated by taking the mean of the MLEs as described above. Due to the presence of short sequences mapping to the induced genes, we set a requirement of 10 sites per transcript rather than the usual 5 to remove these. We quantified abundance using *IsoQuant v3.0.0* and, for ease of viewing, calculated transcript per thousand values by taking the transcript per million normalised output files and dividing the values by 1,000. The clustering and heatmap generation were performed using the *pheatmap* v1.0.12 package.

##### Monocyte LPS stimulation analysis

We plotted the log2FC in both age and counts for each replicate relative to the mean value of the 0h replicates as points with a smoothed line (LOESS method) with the shaded area denoting the 95% confidence intervals.

For the clustering analysis, genes were clustered by differential age relative to the 0H timepoint, thus generating 10 clusters (Table S3). To display the data, four clusters of interest are shown in Fig. 4. The centroids of these 4 clusters were calculated and the 10 genes with the shortest Euclidean distance to the centroid are shown. Cluster robustness was tested by non-parametric bootstrap (Method), and after filtering on robustness, we retained 4 out of 10 initial clusters. In each iteration of the bootstrap, resampled the entire dataset and repeated clustering on the resampled dataset. We compared each original cluster to the most similar bootstrapped cluster, respectively, and calculated the similarity. Clusters which retained their structure after bootstrapping were considered robust (Methods). After filtering on robustness, we retained 4 out of 10 clusters (Figure 4).

For the enrichment analysis, G:profiler (*41*) was used to identify miRNAs and transcription factors (TFs) enriched within each cluster. A list of all genes with >5 reads in the monocyte RNA-seq data was used as the background. Multiple testing correction was performed using the g:SCS threshold (*41*).

##### HEK293 Single cell analysis

Cells containing less than 1000 expressed genes or less than 20 genes that contained calibration fit sites were excluded. Cells were ordered based on the ratio of expression of S-genes/M-genes as defined by Tirosh et al. (*42*). The top 15 S- and M-phase cells were utilised in the subsequent analysis. We focused our analysis on the 554 genes with at least 5 calibration fit sites in the 3’UTR (189,707 total fitted transcripts). Gene and transcript age likelihoods were calculated as described above.

To explore whether transcripts for genes might be expressed cyclically with the cell cycle, we first derived the relative density of transcripts for a given gene for the top S- and M-phase cells, exploring each 2-hour time bin from 0 to 16 hours. For each time bin, we then subtracted the M-phase density from the S-phase density to derive the delta-S/M-density-per-bin. We focused our analysis on the 34 genes that had information in 90% of bins, and with more than 500 total reads across all included cells. Using the non-linear least squares function from the stats package, we fit a sin curve to the delta-densities of the equation Delta = A * sin(B * (Age - C)) + D, with the following starting conditions: A = 0.05, B = 0.6, C = 0 or 6 if convergence not attained, and D = 0. An upper limit was set for B = 1.57, correlating with a minimum cell cycling rate of 4 hours (Tables S8 and S9). For the 25 genes where the model converged within the parameters above, the process was repeated through 750 bootstrapping loops, where in each iteration the S- or M-phase label of a cell was randomly shuffled, with metrics derived including residual sum of squares and a p-value for the B metric. A p-value was derived for the residual-sum-of-squares as the count of bootstrapped results lower than the true value, divided by the total number of bootstrapped results. The period was calculated using Period = 2π/B.

##### Single cell clustering analysis

To cluster the genes, we utilised a biclustering approach suitable for clustering sparse data (*43*, *44*). This approach partitioned the genes by minimising the total sum of squared errors (SSE) between genes in a cluster. This process is repeated through 1000 iterations, comparing iterations via the Rand index to minimise the SSE. Using the mean age per gene we defined 6 clusters, first using MLEs from all transcripts in the top 15 S and M phase cells, and then with transcripts filtered to those less than 4 hours old.

To compare the clustering when transcripts were filtered to those less than 4 hours old, we explored the clustering location of the 3 of 98 canonical cell cycle genes present in the data, *HELLS*, *MDM2*, and *TMPO*. We explored the functions of the 2 clusters containing these genes using the g:Profiler against a background of all the genes expressed in the data, using the tailored g:SCS (g:Profiler: Set, Counts and Sizes) method to correct for multiple testing. Out of the significant results (Table S10), we plotted the gene ontology terms and the most significant miRNA and transcription factor (Fig. S-7).

## Supplementary Tables

Table S1: differentially expressed genes at each timepoint of the human monocyte experiment.

Table S2: differentially aged (6H vs 0H) genes which are not detected by differential abundance in the human monocyte experiment.

Table S3: enriched miRNAs and transcription factors associated with clusters in the human monocyte experiment.

Table S4: plasmids used in this paper.

Table S5: primers, ultramers and g-blocks used in this paper.

Table S6: thermal cycling conditions for reverse transcription and PCR steps of the Nanopore RNA-seq library preparation.

Table S7: Results of fitting sinusoidal curve to differential age in S verse M phase cells.

Table S8: Mean gene ages in S-phase and M-phase cells, with assigned clusters.

Table S9: Mean gene ages in S-phase and M-phase cells with transcripts filtered to those <4 hours, with assigned clusters

Table S10: g:Profiler results for genes in clusters 2 and 4 of the single cell experiment.

## Supplementary Figures

**Fig. S-1.**
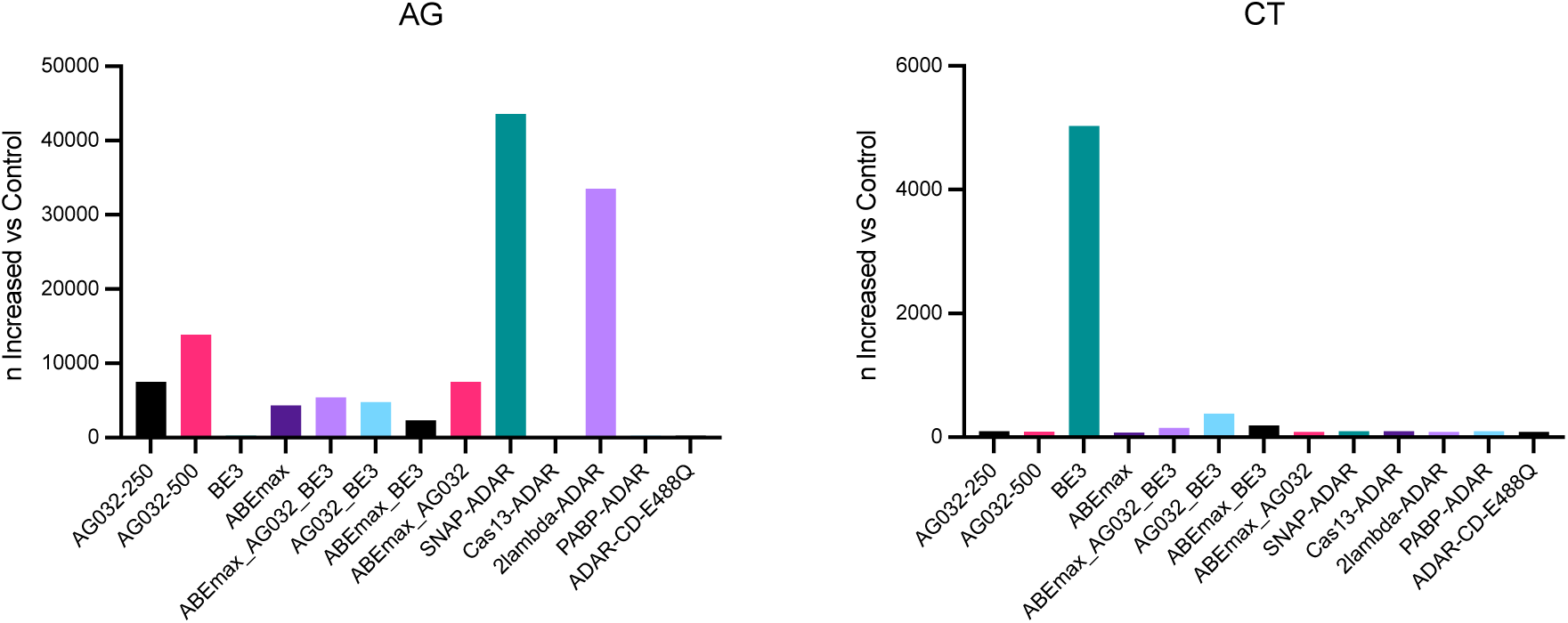
Comparison of first-generation transcriptome-wide, hyperactive editors. Shown are counts of A-to-G (left) and C-to-T transitions (right) detected by JACUSA2 in differential mode relative to the control (empty plasmid). AG032 = ADAR2(E488Q); AG032-250 = 250ug per well; ADAR-CD-E488Q = ADAR2(E488Q) catalytic domain only; ABEmax_AG032_BE3 is an equimolar mixture of three plasmids. Other plasmids are described in Materials and Methods.

**Fig. S-2.**
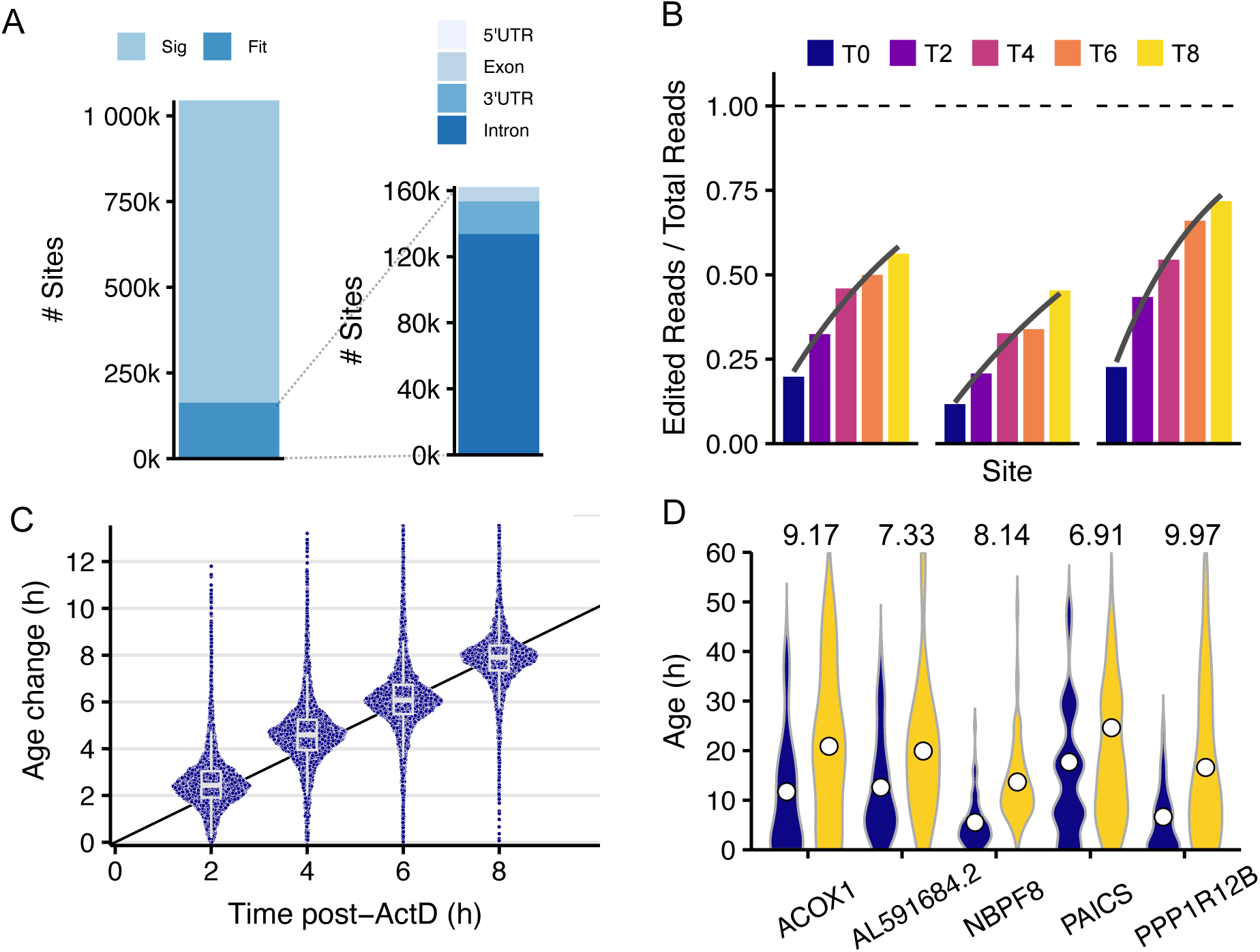
Prediction of transcript ages from endogenous ADAR-mediated A-to-I editing in human cortical neuron culture. (**A**) Left: 1,044,970 adenosines are significantly differentially edited in hiPSC-derived cortical neurons 8 hours after adding actinomycin D. Right: Of those editing sites, 162,741 are well fit by an exponential CDF, and their location within genes are shown as stacked bars. (**B**) Editing fractions at three sites in the 3’UTR of the EIF3M gene are shown at 2 hour increasing time intervals following ActD treatment. The fit of an exponential CDF is shown in grey. (**C**) The mean age of each gene is calculated from the short-read HEK calibration data and the time change compared to the T0 time point (plotted as purple points). A black line of gradient 1 is shown as a guide. Boxplots show the 25th, 50th and 75th quantiles, whiskers extend to the value at most 1.5 * IQR from the respective hinge. (**D**) MLE distributions are shown as violins for the T0 (navy) and T8 (yellow) calibration time points. The mean age is shown as white circles with the Δ age written above.

**Fig S-3.**
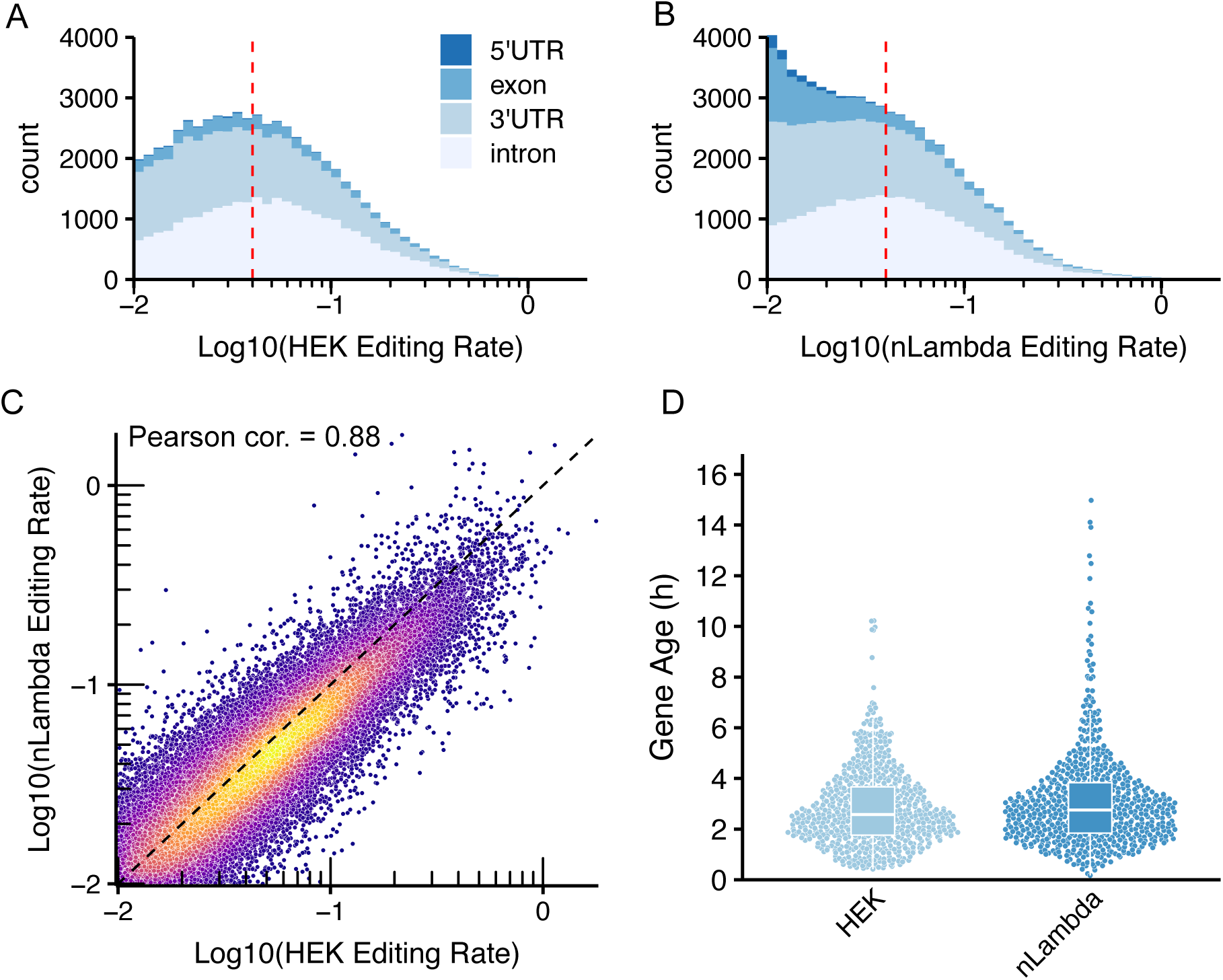
ADAR2(E488Q)-Nlambda contributes extra sites, but these are edited at a low rate and do not contribute useful Timestamps after filtering. **(A)** Stacked histograms showing the distribution of log transformed editing rates of fit sites, broken out by location for the Endogenous HEK (A, n=60,952 sites) and ADAR2(E488Q)-Nlambda expressing HEK (B, n = 68,599) calibrations. A red line is drawn at editing rate = 0.04 h^−1^, above which the two distributions highly correlate. (**C**) The log-transformed editing rates of shared fit sites (n=36,768) between endogenous and ADAR2(E488Q)-Nlambda strongly correlate (Pearson correlation coefficient = 0.88). (**D**) Gene ages are estimated by T2 from the T0 time point of the Endogenous HEK calibration experiment using either the endogenous HEK or the nLambda HEK calibration datasets. The fit sites are filtered to 3’ UTR sites in genes that have at least five 3’UTR sites (HEK n = 26,463, nLambda n = 26,302 sites).

**Fig. S-4.**
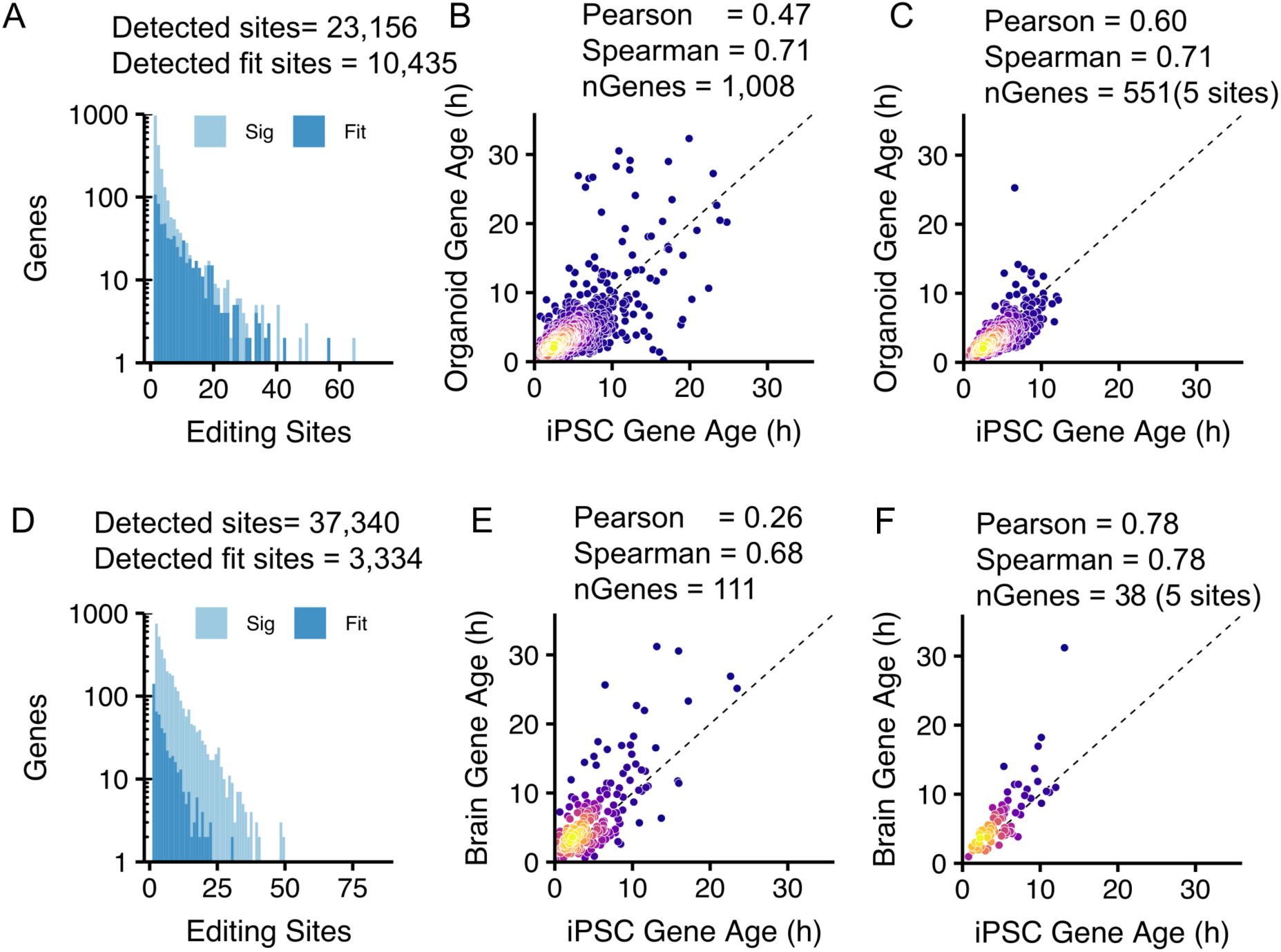
Sequencing data from human intestinal organoids (A-C) and human brain slices (D-F). (A and D) The distribution of editing sites across the transcriptome in human intestinal organoids and brain slices, respectively. (B and E) Correlation of mean ages of genes with at least 1 fit site from the cortex calibration. (C and F) Correlation of mean ages of genes with at least 5 fit sites from the cortex calibration.

**Fig, S-5.**
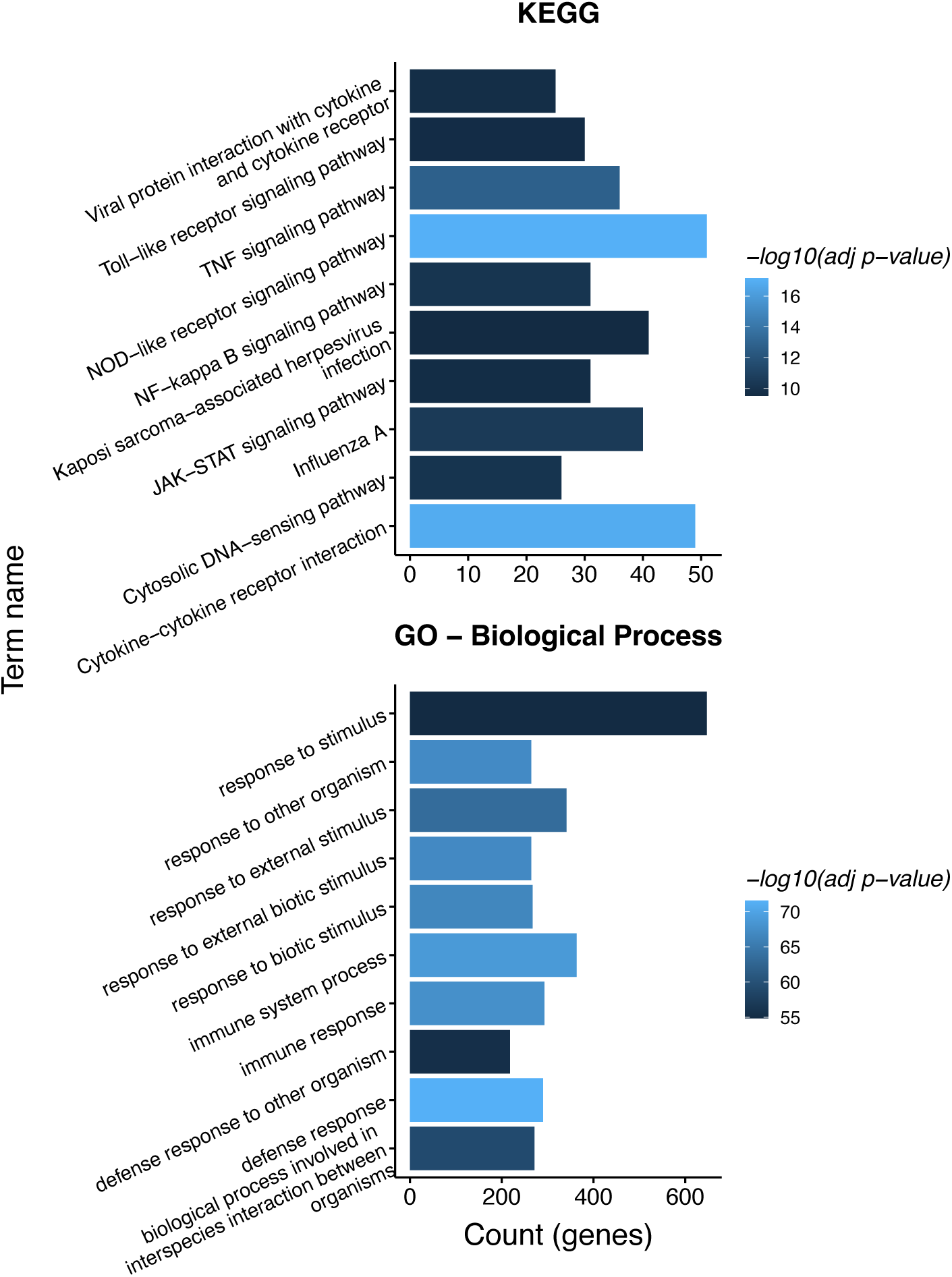
Human CD14+ monocytes were stimulated with bacterial LPS and then lysed at intervals. (A) Gene ontology enrichment analysis, performed using ClusterProfiler (*45*, *46*), indicates activation of canonical M1 Macrophage activation pathways.

**Fig. S-6.**
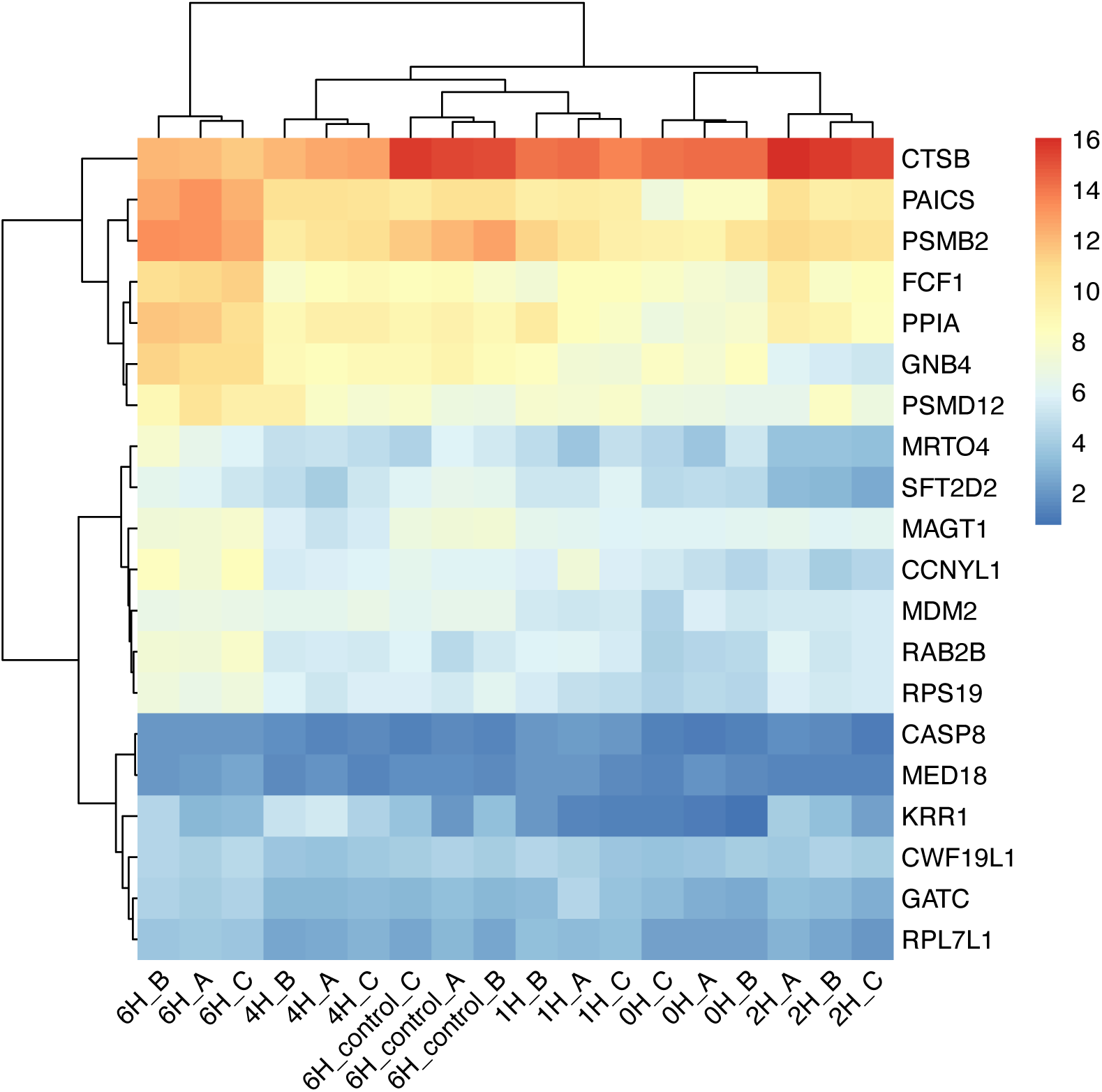
Heatmap showing genes with a significant change in mean age (6H vs 0H), clustered by gene and replicate. Replicates cluster together.

**Fig. S-7.**
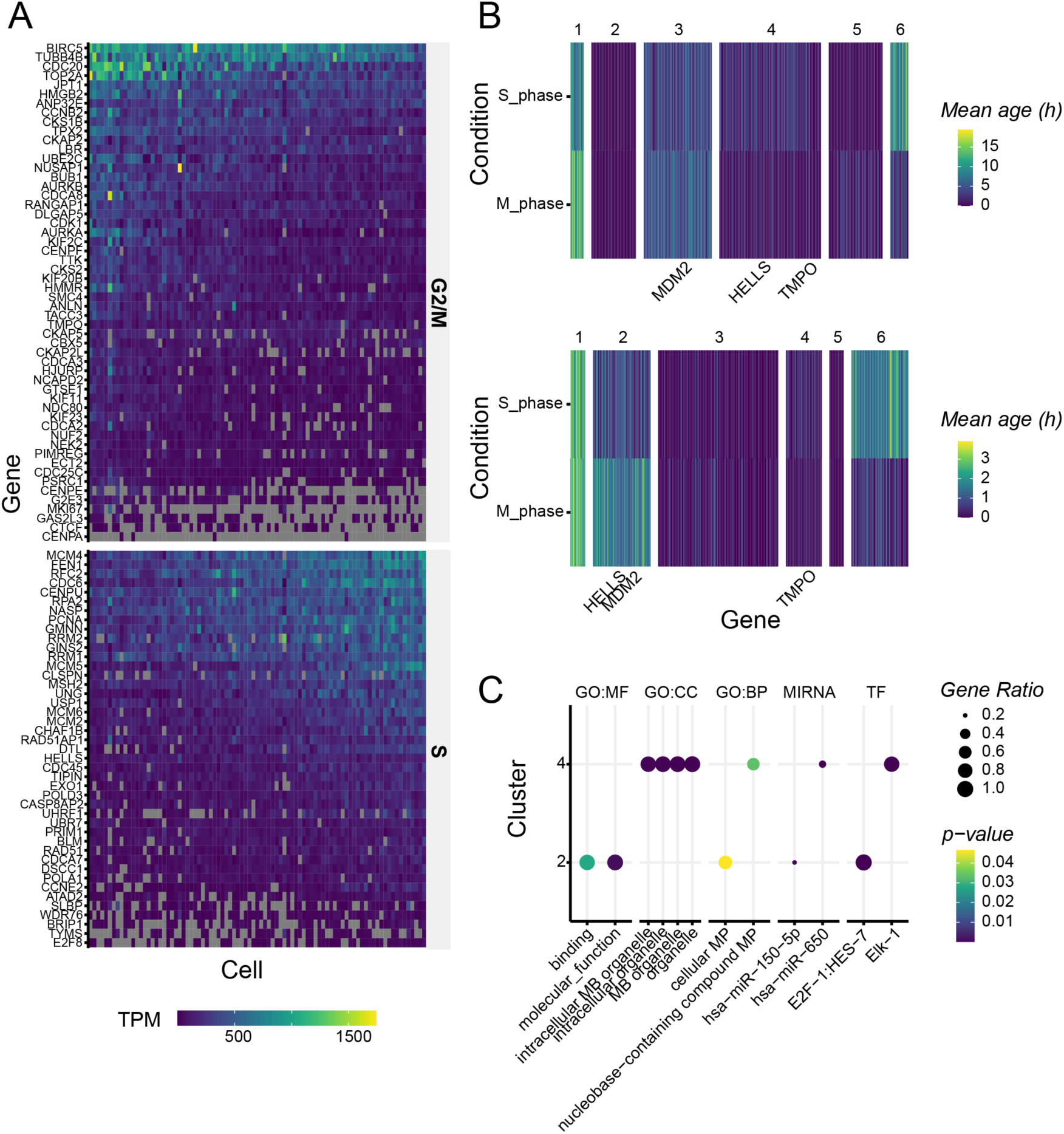
(**A**) Gene expression of HEK293 single cell analysis, showing the expression of G2/M phase genes (top), and G1/S phase genes (bottom), ordered by the ratio of mean G2/M verse S. Cells were ordered via mean ratio of expression of M/S phase genes, allowing selection of the top 15 G1/S phase cells and the top 15 S/M phase cells. (**B**) Results of a clustering analysis using a biclustering approach specific for sparse data (*43*) Clustering of genes in S phase cells verse M phase cells, unfiltered for transcript age (top). Note that *HELLS* and *TMPO* cluster together in cluster 4, suggesting a ‘cell cycle cluster’ that does not differentiate between S and M phase. This is improved when transcripts are restricted to those <4 hours old (bottom) where *HELLS* clusters with *MDM2* in cluster 2 (younger in S phase), while *TMPO* clusters in cluster 4 (younger in M phase). (**C**) Results of g:Profiler (*41*) enrichment analysis for the *HELLS* containing cluster 2 (bottom) and *TMPO* containing cluster 4 (top), showing all gene ontology results and most significant results from miRNA and transcription factor analysis. Cluster 2 was associated with DNA binding as expected in the S-phase, as well as the E2F transcription factor which has been associated with progression from G1 to S phase (*47*, *48*) long with the highlighted microRNA hsa-miR-150 (*49*). The cluster containing TMPO, a mitosis related gene, was enriched for membrane bound organelles, as well as the hsa-miR-650 which has been associated with the metaphase of mitosis (*49*) and the transcription factor Elk-1, which localises to the mitotic spindles during the cell cycle (*50*).

**Fig S-8.**
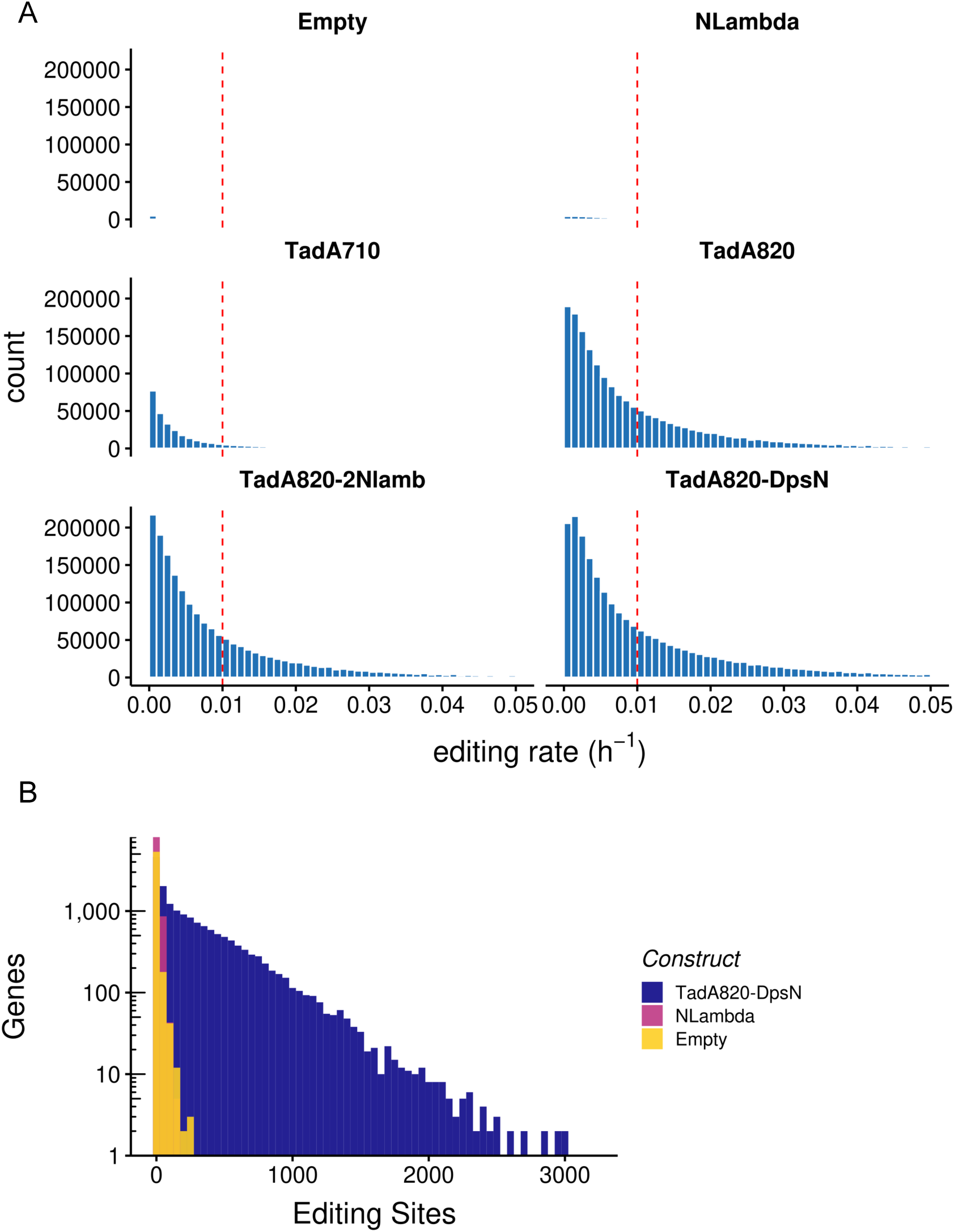
Second-generation hyperactive editors developed using E.coli tRNA-specific adenosine deaminase (TadA). The editing rates and distribution of editing sites of the second-generation editors acting on the human transcriptome are shown. (**A**) A calibration experiment consisting of two timepoints (0 and 8 hours after actinomycin D) was performed. A linear model was fit to the numbers of edits at detected sites to approximate the editing rate lambda from Equation 1 (Main). Histograms of the editing rates are shown. (**B**) Histogram showing the transcriptome-wide distribution of editing sites identified by JACUSA2 for the empty vector control (endogenous editing), Nlambda-ADAR2 and TadA8.20-Dps-N.

